# Stereological characterization of neurogenic niche secretory cell types in the mouse dorsal dentate gyrus

**DOI:** 10.1101/847350

**Authors:** Joshua D. Rieskamp, Patricia Sarchet, Bryon M. Smith, Elizabeth D. Kirby

## Abstract

Adult neurogenesis in the dorsal dentate gyrus (DG) subregion of the mammalian hippocampus supports critical cognitive processes related to memory. Local DG cell populations form a neurogenic niche specialized to regulate adult neurogenesis. Recently, DG astrocytes, microglia, endothelia, and neural stem cells have been identified as sources of neurogenesis-modulating secreted factors. Accurately estimating the size of these cell populations is useful for elucidating their relative contributions to niche physiology. Previous studies have characterized these cell types individually, but to our knowledge no comprehensive study of all these cell types exists. This is problematic because considerable variability in reported population size complicates comparisons across studies. Here, we apply consistent stereological methods within a single study to estimate cell density for neurogenesis-modulating secretory cell types in the dorsal DG of adult mice. We used immunohistochemical phenotypic markers to quantify cell density and found that stellate astrocytes were the most numerous followed by endothelia, intermediate progenitors, microglia, and neural stem cells. We did not observe any significant sex differences in cell density. We expect our data will facilitate efforts to elucidate the role of DG secretory cell populations in regulating adult neurogenesis.

## Introduction

The mammalian hippocampus is well studied for its role in episodic memory, spatial navigation, and mood regulation (1). Within the hippocampal circuit, the dorsal dentate gyrus (DG) subregion mediates crucial computational processes related to memory, such as pattern separation (2,3). These proposed functional roles likely relate to the distinct neuroanatomical features of the DG, including its organization along the longitudinal (dorsal-ventral or septal-temporal) axis (1).

The DG is unique from most adult brain circuits because it hosts a specialized niche where neural stem cells (NSCs) continuously generate granule neurons throughout life. Adult hippocampal neurogenesis (AHN) occurs in most mammals, possibly including humans (4–7). Rodent studies implicate new granule neurons generated via adult neurogenesis in several key cognitive tasks carried out by the DG (8). Furthermore, experimental manipulations that mitigate age-related declines in neurogenesis improve cognitive performance (9,10), raising the possibility that targeting neurogenesis could be therapeutically useful. Therefore, developing a comprehensive picture of the factors that regulate adult hippocampal neurogenesis is a topic of intense investigation.

AHN relies on external regulation from the niche microenvironment. Recent studies demonstrate that secreted factors modulate key facets of neurogenesis such as NSC proliferation dynamics and adult-born neuron survival/maturation (11). These secreted factors originate from multiple niche-resident cell types including vascular endothelia cells, astrocytes, microglia, and even undifferentiated NSCs (12–16).

As these multiple cell types contribute to the neurogenesis-modulating secretome, it is useful to know their relative numbers. However, these cell types have not been subjected to the same level of stereological characterization as other DG populations like mature and immature granule neurons. Furthermore, most stereological studies examine a select few cell types, and while efforts have been made to generate comprehensive atlases (17), these databases currently lack information about essential DG secretory cell types such as NSCs and vascular endothelial cells. Therefore, one must combine multiple sources to obtain a comprehensive estimate of DG secretory cell composition. This approach is problematic because numbers vary substantially across studies, even for commonly quantified cell types (18), leading to considerably different conclusions depending on which data is used to draw comparisons.

To circumvent this issue, here we apply consistent stereological methods within the same study to estimate the relative numerical densities of NSCs, astrocytes, microglia, and vascular endothelial cells in the DG. To our knowledge, this represents the first effort to quantify these AHN-modulating secretory cell types within a single study.

## Results

We used immunohistochemistry to identify cells expressing phenotypic markers of neurogenesis-modulating secretory cell populations within the DG: NSCs and their intermediate progenitor cell progeny (IPCs), mature stellate astrocytes, microglia, and vascular endothelial cells. Cell density estimates were derived for adult wildtype C57BL/6 mice using the optical fractionator method to count cells within a region of interest (ROI) encompassing all layers of the DG (molecular, granular, and hilar) at the dorsal (septal/rostral) pole of the dorsal-ventral axis (Figure 1A-B).

**FIGURE 1.**
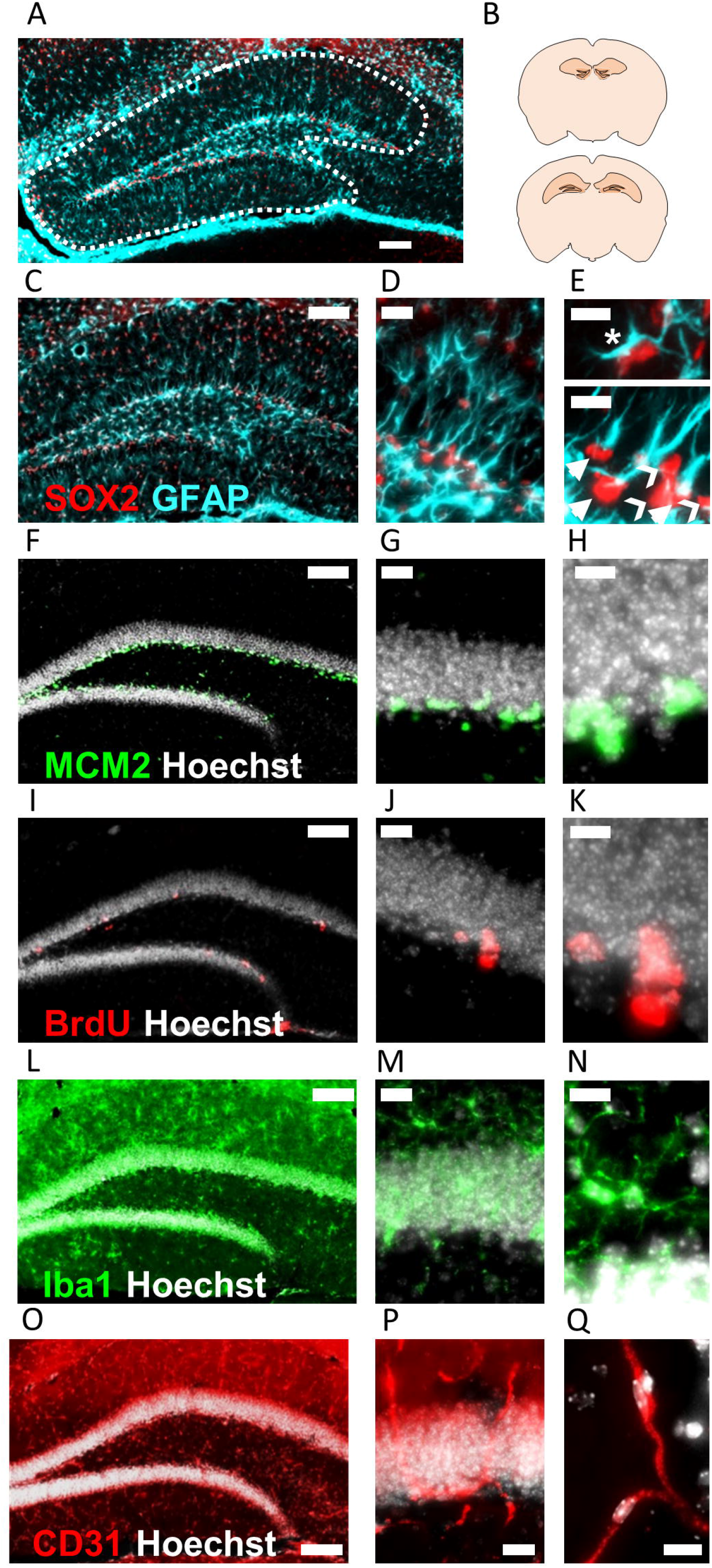
Representative immunolabeling of neural stem and progenitor cell, astroglial, microglial, endothelial, and proliferative cell markers in the dorsal dentate gyrus (DG). (**A-B**) The ROI used for cell counts encompassed all layers of the DG and spanned the septal (dorsal) pole of the septal-temporal axis. (**C-E**) Representative SOX2 and GFAP immunolabeling. Mature astrocytes had SOX2+ nuclei, GFAP+ cytoplasm, and stellate morphology (**E**, **top, asterisk**). Radial glia like neural stem cells had SOX2+ nuclei in the SGZ, GFAP+ cytoplasm, and a radial process spanning the GCL (**E, bottom, arrows**). Neural intermediate progenitors had SOX2+ nuclei in the SGZ lacking GFAP+ cytoplasm (**E**, **bottom, chevrons**). (**F-H**) MCM2 immunolabeling and (**I-K**) BrdU immunolabeling 2 hours post injection to reveal actively cycling cells. (**L-N**) Representative Iba1 immunolabeling for microglia. (**O-Q**) CD31 immunolabeling to detect vascular endothelial cells. Scale bars represent 100 μm (**A, C, F, I, L, O**), 20 μm (**D, G, J, M, P**), or 10 μm (**E, H, K, N, Q**).

### NSCs and astrocytes

Many of the protein markers expressed by adult NSCs are also present in mature astrocytes (19). Therefore, we distinguished these cell populations based on a combination of SOX2 and GFAP immunolabeling along with cell morphology (20). We identified radial glial-like neural stem cells (RGLs) as having SOX2+ nuclei located in the subgranular zone (SGZ) and GFAP+ cytoplasm with a radial process extending into the granule cell layer (Figure 1D-E). The progeny of RGLs, neural intermediate progenitor cells (IPCs), were identified as SOX2+ nuclei located in the SGZ that lacked GFAP+ cytoplasm (Figure 1D-E). Mature protoplasmic astrocytes had SOX2+ nuclei located in any region of the DG and GFAP+ cytoplasm with stellate processes (Figure 1D-E). We found that mature astrocytes were the most abundant of these three populations (8274.7 ± 1109.7, 9232.4 ± 1095.1 cells/mm3, females and males, respectively) followed by IPCs (4532.2 ± 469.7, 5220.1 ± 820.2 cells/mm3), then RGLs (1364.9 ± 132.0, 1228.6 ± 59.5 cells/mm3, Figure 2A). Males and females did not significantly differ in number of astrocytes (p>0.05, t=0.61), RGLs (p>0.05, t=1.02), or IPCs (p>0.05, t=0.68).

**FIGURE 2.**
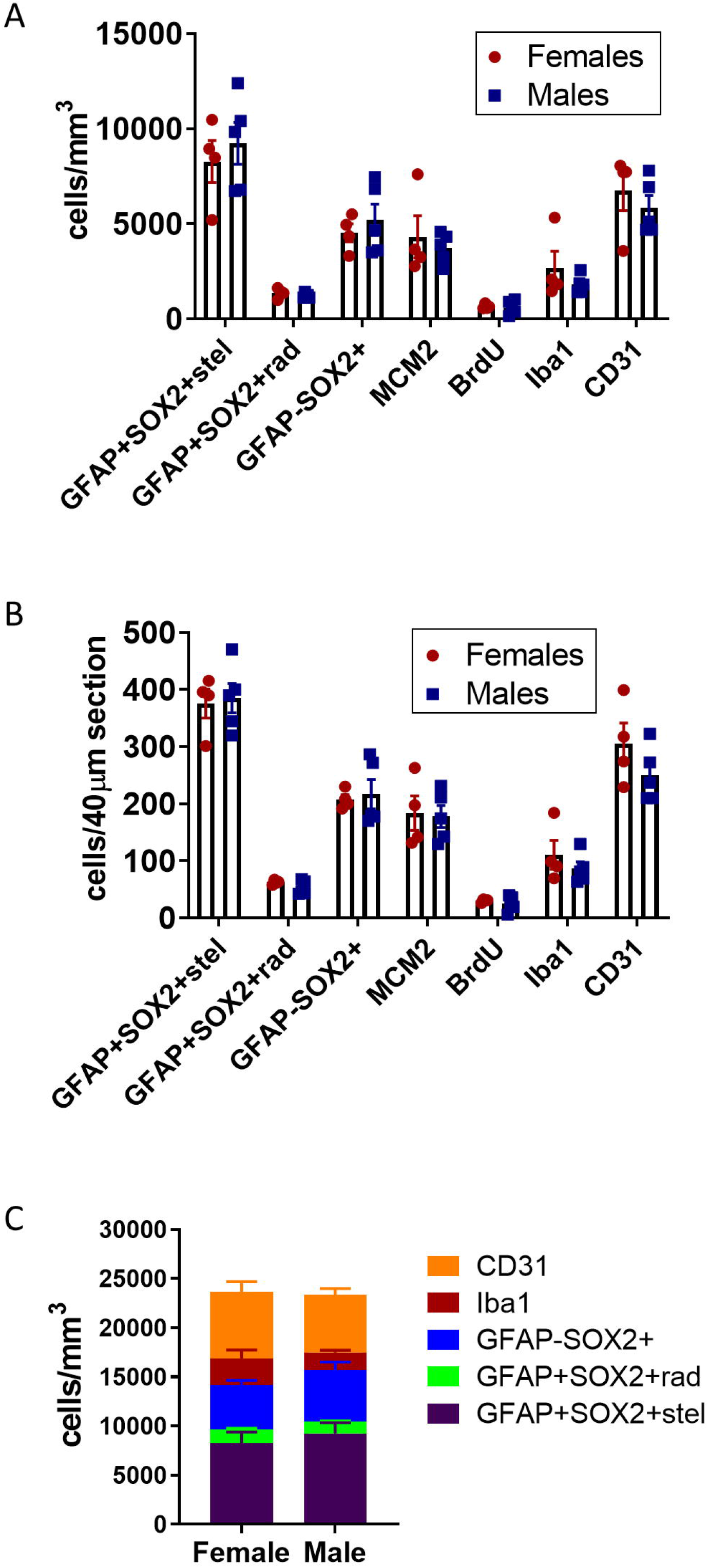
Stereological estimates of all cell types surveyed in the current study. Data are represented as cell density (**A**) and number of cells in a 40 μm slice (**B**). (**C**) Cell density estimates of mutually exclusive cell type categories (for example, MCM2+ cells might overlap with multiple other cell types and is thus not included). Data is mean ± SEM from male (n=5) and female (n=4) adult mice. No significant sex difference for any cell type was detected via unpaired, two-tailed t-test (p>0.05).

### Mitotically active cells

To expand the quantification of progenitor cells, we used two markers of cellular mitosis. The adult DG contains various populations of mitotically active cells, which can be identified using endogenous cell cycle markers such as MCM2 or the exogenous S phase maker bromodeoxyuridine (BrdU) (21,22). Previous studies have shown that the majority of MCM2+ or recently BrdU-labeled cells in the DG are IPCs located within the SGZ, with a sparser number of proliferative glial cells located outside the SGZ (23,24). Consistent with these previous findings, we found MCM2+ cells throughout the entire DG, but the greatest density was found within the SGZ (Figure 1F-H). To obtain counts most directly related to IPC number, only MCM2+ cells in the SGZ were quantified. The density of SGZ MCM2+ cells did not significantly differ between females and males (4324.1 ± 1106.9 vs 3740.7 ± 356.8 cells/mm3, p>0.05, t=0.55) (Figure 2A). We obtained similar results quantifying BrdU+ cells after a single BrdU injection administered 2 hours before perfusion. BrdU labeled cells were found throughout the entire extent of the DG, but most commonly within the SGZ (Figure 1I-K). To obtain counts most directly related to the number of IPCs in S-phase, only BrdU+ cells in the SGZ were quantified. The density of SGZ BrdU+ cells was not different between females and males (658.4 ± 61.9 vs 597.3 ± 162.1 cells/mm3, p>0.05, t=0.32) and represented a smaller subset of cells than all cycling (i.e. MCM2+) cells, as expected (Figure 2A).

### Microglia

Iba1 + microglia were distributed throughout every region of the DG (Figure 1L-N) and were similar in number to RGLs. No sex difference was found in Iba1 + cell density (2677.1 ± 892.1 vs 1816.8 ± 210.2 cells/mm3, p>0.05, t=1.05, Figure 2A).

### Vascular endothelial cells

Vascular endothelial cells within the DG were identified by CD31 immunolabeling and formed vascular networks throughout all subregions (Figure 1O-Q). The density of CD31+ cells was intermediate between the density of astrocytes and IPCs, with no sex difference (6768.7 ± 1067.3 vs 5849.0 ± 639.4 cells/mm3, p>0.05, t=0.78, Figure 2A).

### Summary

Cell density estimates for each cell marker quantified in the current study are depicted in Figure 2A. To facilitate comparisons with counts performed in sections cut to a commonly used thickness for free-floating immunohistochemistry, data is also presented as estimates of cell number in a 40 μm slice (Figure 2B). In addition, we provide an estimate of relative population sizes for the major cell types quantified using mutually exclusive markers for RGLs, IPCs, astrocytes, microglia, and endothelial cells (Figure 2C). This presentation of the data allows an appreciation of the approximate relative population sizes of the major DG secretory cell types surveyed in this study.

## Discussion

Adult hippocampal neurogenesis in rodents occurs within the DG, a niche comprised of neurogenesis-supporting cells. So far, studies have identified several cell types – astrocytes, microglia, endothelial cells, and neural stem cells – which regulate neurogenesis via secreted factors (12–16). Here, we quantified these cell types using stereology to obtain an estimate of the relative population densities of neurogenesis-modulating secretory cells within the dorsal DG.

We found that GFAP+SOX2+ stellate astrocytes were the largest cell population in our study, followed closely by CD31+ vascular endothelial cells. Next most abundant were GFAP-SOX2+ IPCs, Iba1 + microglia, and GFAP+SOX2+ RGLs. Notably, we compared densities obtained from both male and female mice for each cell type and found no significant sex differences. While a previous study reports sex differences in DG astrocyte and microglia density (Mouton et al., 2002), quantification for males and females were obtained from separate rounds of tissue processing and counting. Though the authors made efforts to maintain consistent methodology, small but systematic differences between cohorts could influence the results. In our study, where both sexes were processed and counted in a single cohort, we did not observe sex differences in astrocytes, microglia, or any other cell type surveyed. To our knowledge, this is the first study to quantify this set of cell types in the DG of both sexes by applying the same methodology to a single cohort of mice.

Previous studies have estimated population size for each of the DG cell types studied here individually. However, comparison of our data with previous work is complicated by the variability in population estimates across studies (18). For example, estimates of astrocyte density in the DG range from 19,900 cells/mm^3^ (25) to 55,700 cells/mm^3^ (26). Despite these studies using unbiased stereology, variability could arise for biological (different cohorts of mice), methodological (different markers and anatomical boundaries), and computational (different extrapolation from sampled counts to total population density estimates) reasons. Even small differences in sampling parameters and adjustments for tissue shrinkage become amplified when calculating estimates of total population size and tissue volume (18). How the data derived from differing methods can be adjusted for comparison is not immediately clear. This variability combined with the tendency of most studies to examine a select few cell types complicates attempts to compare cell population sizes across studies.

To circumvent these issues, we sought to use consistent methodology to obtain estimates for a comprehensive set of DG cell types previously implicated in modulating neurogenesis via secreted factors. All data in this study were generated using tissue from a single cohort of male and female mice of the C57BL/6 strain, one of the most commonly used rodent species in biosciences research (27). Therefore, the main advantage of this dataset is the validity of internal comparisons between various cell types in a common research model species. For example, when drawing comparisons based on data reported in separate studies, the microglia to astrocyte ratio could range from approximately 1:1 (25,26) to 1:6 (28,29). By comparison, using our estimates, we report a microglia to astrocyte ratio of roughly 1:4, similar to the approximately 1:3 ratio reported by others when both cell types were quantified within the same study (26,29). This test case highlights the utility of counts obtained within a single study for making accurate comparisons of relative population sizes.

Important limitations to this dataset should be noted. Counts were performed exclusively in the dorsal (rostral/septal) DG with no separate delineation of the layers (i.e. molecular, granular, hilar) or inclusion of ventral DG populations. We chose to focus on dorsal DG because it represents a functional unit with known roles in mediating memory functions related to pattern separation and temporal encoding (3). While the degree to which the DG dorsal-ventral axis constitutes discrete brain regions versus overlapping functional gradients is a matter of debate (1), distinct gene expression, connectivity, and behavioral correlates at the dorsal and ventral poles (30) provide justification for separate analysis. Given reports of disparate cell densities along the DG longitudinal axis (31), future studies should quantify the major secretory cell populations of ventral DG. Additionally, similar comprehensive analysis of neurogenesis-modulating cell types in the subventricular zone and hypothalamus could elucidate regionspecific mechanisms for regulating adult neurogenesis.

Overall, we have provided stereological density estimates for neurogenesis-modulating secretory cell types of the dorsal DG. Our data facilitates direct comparisons of multiple cell types, which we expect will be useful for refining hypotheses about the relative contributions of these cell types to regulating adult neurogenesis.

## Methods

### Animals

All animal use was in accordance with institutional guidelines approved by the Ohio State University Institutional Animal Care and Use Committee and with recommendations of the National Institutes of Health Guide for the Care and Use of Laboratory Animals. C57BL/6 mice (5 male, 4 female) were acquired from Jackson Laboratories at 7 weeks of age and housed in a vivarium at Ohio State University with 24h ad libitum water and standard mouse chow on a 12h light-dark cycle for 1 week. Mice received one injection of 150 mg/kg bromodeoxyuridine (Sigma) dissolved in physiological saline, and 2h later were anesthetized with 87.5 mg/kg ketamine/12.5 mg/kg xylazine and transcardially perfused with 0.1 M phosphate buffered saline (PBS).

### Tissue processing

Brains were fixed for 24h in 4% paraformaldehyde in 0.1 M phosphate buffer then equilibrated for at least 2 days in 30% sucrose in PBS, both at +4°C. They were then sliced on a freezing microtome (Leica) in a 1 in 12 series of 40 μm slices. Slices were stored in cryoprotectant at −20°C. Immunohistochemical staining was performed on free-floating sections as previously described (32). Briefly, sections were rinsed three times in PBS, incubated in blocking solution (1% normal donkey serum, 0.3% triton-X 100 in PBS) for 30 min then incubated in primary antibody diluted in blocking buffer overnight at 4°C on rotation. The next day, sections were rinsed and incubated in secondary antibody in blocking solution at room temperature for 2 hours, followed by 10 min in Hoechst 33342 (Invitrogen) diluted 1:2000 in PBS. After rinsing, sections were mounted on superfrost plus glass slides (Fisher) and cover-slipped with Prolong Gold Anti-fade medium (Invitrogen). After drying, slides were stored in the dark at 4°C until imaging. For BrdU immunohistochemical labeling, the above procedures were followed with the addition of a 30 min incubation in 2N HCl at 37°C to denature DNA before the blocking step.

### Stereological cell counts

Stereological cell counts were performed in a single series of every 12th slice for each cell phenotype marker. Images of the dorsal DG were captured using Zeiss AxioImager.M2 microscope and a Zeiss MRc digital camera. Cells were counted at 40x magnification with oil immersion using the optical fractionator method (StereoInvestigator). The counting frame had an area of 10,000 μm^2^ and a height of 15 μm with 5 μm guard zones. Proliferating cells were identified using the nuclear markers in **Table 1**. Endothelial cells and microglia were identified by the cytoplasmic markers in **Table 1** surrounding a Hoechst+ nucleus. Radial glia-like neural stem cells (RGLs) were identified as SOX2+ nuclei in the SGZ surrounded by GFAP+ cytoplasm with an apical process extending into the granular cell layer. Neural intermediate progenitor cells (IPCs) were identified as SOX2+ nuclei in the SGZ lacking cytoplasmic GFAP. Astrocytes were identified as SOX2+ cells with GFAP+ stellate processes located in any DG subregion. The mean number of cells counted for each marker are listed in **Table 2**.

**Table 1.** Antibodies used for immunohistochemical identification of cell phenotype. All antibodies were validated in previous work or compared to an appropriate control to ensure specificity of immunolabeling.

| Primary antibodies |  |  |  |  |  |  |  |
| --- | --- | --- | --- | --- | --- | --- | --- |
| Antibody | Host species | Cell type | Antigen retrieval | Vendor | Product # | Dilution | Ref/control |
| $\alpha$ Sox2 | rat | RGL, IPC, astrocyte | None | eBioscience | 14-9811 | 1:1000 | (33) |
| $\alpha$ GFAP | rabbit | RGL, astrocyte | None | Dako | Z-0334 | 1:1000 | (34) |
| $\alpha$ MCM2 | rabbit | Proliferating cells | None | Cell Signaling | 4007 | 1:500 | (35) |
| $\alpha$ BrdU | mouse | Proliferating cells | 2N HCl, 37°C | BD Biosciences | BDB 347580 | 1:500 | No BrdU injection |
| $\alpha$ Iba1 | goat | Microglia | None | Abcam | Ab5076 | 1:2000 | (32) |
| $\alpha$ CD31 | rat | Endothelia | None | BD Pharmingen | 550274 | 1:100 | (36) |
| **Secondary antibodies** |  |  |  |  |  |  |  |
| 647 $\alpha$ rabbit | donkey | GFAP, MCM2 | N/A | Invitrogen | 1:500 | A31573 | No primary |
| 594 $\alpha$ rat | donkey | Sox2, CD31 | N/A | Invitrogen | 1:500 | A21209 | No primary |
| 488 $\alpha$ mouse | donkey | BrdU | N/A | Invitrogen | 1:500 | A21202 | No primary |
| 488 $\alpha$ goat | donkey | Iba1 | N/A | Invitrogen | 1:500 | A11055 | No primary |

**Table 2.** Total counts, distribution, error estimators, measured tissue thickness, and number of sections sampled for all cell types surveyed in the current study. For each cell type, the total cells counted and percentage of cells counted in the middle 5 consecutive 1 μm z-sections and top 5 consecutive 1 μm z-sections immediately below the guard zone are listed (mean ± SEM). Error estimators are listed as Gunderson coefficients (with smoothness *m* = 0 or *m* = 1). For each round of immunostaining, the measured thickness (mean ± SEM) and number of sections samples is provided. Thickness was manually measured in 3 locations per slice to adjust for shrinkage relative to the 40 μm starting thickness.

| Cell type marker | Total Cells Counted<br>(Mean ± SEM) | Distribution<br>(% of all counted cells, mean ± SEM) |  | Gundersen-Jensen CE *m* = 0 | Gundersen-Jensen CE *m* = 1 |
| --- | --- | --- | --- | --- | --- |
|  |  | Top 5 μm | Middle 5 μm |  |  |
| GFAP+SOX2+stel | 308.1 ± 7.5 | 32.32 ± 0.95 | 37.55 ± 0.73 | 0.144 | 0.064 |
| GFAP+SOX2+rad | 46.4 ± 4.1 | 34.44 ± 1.75 | 33.48 ± 2.37 | 0.203 | 0.154 |
| GFAP-SOX2+ | 148.3 ± 8.2 | 30.07 ± 2.17 | 36.06 ± 1.83 | 0.164 | 0.088 |
| MCM2 | 140.1 ± 10.5 | 38.47 ± 3.69 | 33.63 ± 2.52 | 0.188 | 0.094 |
| BrdU | 22.7 ± 2.7 | 37.25 ± 5.32 | 20.58 ± 5.46 | 0.287 | 0.243 |
| Iba1 | 74.3 ± 6.6 | 35.49 ± 0.76 | 34.75 ± 1.76 | 0.164 | 0.120 |
| CD31 | 175.1 ± 7.7 | 30.21 ± 1.15 | 39.07 ± 1.97 | 0.161 | 0.083 |

**Table 2 | Total counts, distribution, error estimators, measured tissue thickness, and number of sections sampled for all cell types surveyed in the current study.**
| Immunostaining Round | Measured Tissue Thickness (mean ± SEM) |  | Sections surveyed | Sites surveyed (mean ± SEM) |
| --- | --- | --- | --- | --- |
|  | Males | Females |  |  |
| GFAP, SOX2, BrdU | 25.3 ± 0.09 | 26.6 ± 1.33 | 3 | 56.8 ± 1.3 |
| CD31 | 25.9 ± 0.80 | 24.8 ± 0.75 | 3 | 60.6 ± 1.3 |
| MCM2, Iba1 | 26.3 ± 0.69 | 25.5 ± 0.86 | 3 | 59.7 ± 1.5 |

### Data analysis and statistics

Cell density was calculated from cell count divided by sampled DG tissue volume. Because immunohistochemical processing causes tissue shrinkage, sampled tissue volume was estimated using sampled area adjusted by the proportional change in measured post-processing tissue thickness (i.e. thickness measured in 3 locations per slice using StereoInvestigator versus the sliced thickness of 40 μm). Mean thickness measurements obtained for each round of immunostaining are reported in **Table 2**. For each cell type, cell densities in males and females were compared using an unpaired, two-tailed t-test (Prism GraphPad).

### Cell distribution

Incomplete antibody penetration can interfere with obtaining accurate stereological estimates. We verified complete antibody penetration by comparing the number of cells for each maker counted in 5 consecutive 1 μm z-sections in the middle of the tissue compared to the 5 consecutive 1 μm z-sections immediately below the guard zone (top). We observed similar counts obtained from the middle z-sections compared to the top, suggesting uniform antibody penetration (**Table 2**).

### Error estimates

Previous work suggests that the Gundersen-Jensen coefficient of error (CE) estimator (37) is useful for evaluating the precision of stereological estimates in the hippocampal structure (38). The mean Gundersen-Jensen CE for each cell type is reported in **Table 2**.

## Author Contributions

PS and BMS processed and stained the tissue. EDK performed the stereological counts. JDR and EDK performed statistical analysis. JDR and EDK wrote the manuscript.

## Acknowledgements

The authors would like to thank Dr. Kathryn Lenz for use of equipment.

